# Heat efficiently inactivates coronaviruses inside vehicles

**DOI:** 10.1101/2021.09.08.459486

**Authors:** Krithika P. Karthigeyan, Chloe Flanigan, Denis Jacob Machado, Alper A. Kiziltas, Daniel A. Janies, Jay Chen, David Cooke, Marcia V. Lee, Linda J. Saif, Sonny Henegar, Jeff Jahnes, Deborah F. Mielewski, Jesse J. Kwiek

## Abstract

Heat is an established method to inactivate coronaviruses, and there is utility in using heat to reduce viral load on common touch points in vehicles exposed to a person shedding SARS-CoV-2. As SARS-CoV-2 is a Biosafety level (BSL)-3 pathogen, real world testing of heat as a sanitation method for public and private vehicles becomes a challenge, requiring a surrogate coronavirus that can be handled safely outside of a BSL-3 facility. In this study, we used Bovine Coronavirus (BCoV) as a surrogate for SARS-CoV-2 to test the efficacy of heat-based betacoronavirus inactivation. *In vitro*, a 30-minute exposure to 56°C completely inactivated BCoV in solution, and a 15-minute exposure reduced recovery of BCoV >1000-fold. When heated to 56°C for 15 minutes, the infectivity of BCoV spotted and dried on typical porous and non-porous automobile interior materials was reduced by 99 - 99.99%. When BCoV was spotted and dried on hard plastic (seat) material placed inside an out of service transit bus, 56°C heat for 30 minutes reduced BCoV infectivity 85 - 99.5%. Thus, 56°C is an accessible, rapid, and effective method to inactivate coronaviruses inside motor vehicles.

## Introduction

Severe Acute Respiratory Syndrome related Coronavirus-2 (SARS-CoV-2), the causative agent of Coronavirus disease 2019 (COVID-19), can be inactivated by exposure to heat,^1–5^ like several other viruses.^1,6,7^ Heat sensitivity of viruses such as SARS-CoV-2 can be leveraged to sanitize common touchpoints of public and private vehicles. This sanitization can be done as part of routine cleaning or as a sanitization protocol following exposure of the vehicle to a person shedding viral particles.

Environmental surveillance has highlighted the persistence of SARS-CoV2 particles in dust indoors,^8^ and several studies have documented the prevalence of SARS-CoV-2 particles in ambient air and surfaces of public vehicles.^9,10^ We sought to test the ability of heat to inactivate coronaviruses on high-touch materials commonly found in both cars and public transportation vehicles. Owing to its classification as a Biosafety level (BSL) 3 pathogen, SARS-CoV-2 cannot be tested in field studies, so we elected to use Bovine Coronavirus (BCoV). BCoV has been used as a surrogate for SARS and Middle Eastern Respiratory Syndrome (MERS) coronaviruses,^11^ and like SARS and SARS-CoV-2, BCoV belongs to the *Betacoronavirus* genus of the *Coronaviridae* family. Similar to SARS-CoV-2, BCoV infects the upper and lower respiratory tract and gastrointestinal tract in cattle, and BCoV particles are shed both in fecal secretions as well as upper respiratory tract secretions.^12^ Using BCoV as a surrogate for SARS-CoV-2, we observed that heating materials to a surface temperature of 56°C followed by a hold at 56°C for 15-30 minutes effectively inactivated BCoV on relevant materials, both in a controlled laboratory environment and in situ on seat material placed inside an out-of-service public transportation bus.

## MATERIALS & METHODS

### Cells and Viruses

Madin Darby Bovine Kidney (MDBK) cells were maintained in advanced minimal essential medium (AMEM, Gibco) supplemented with 5% heat-inactivated Fetal Bovine Serum (FBS), 2 mM L-Glutamine (Gibco), and 1% Antibiotic/Actinomycotic cocktail (Gibco).^13^ BCoV-Mebus (GenBank: U00735.2) was used in all assays.^14^ All experiments were approved by the OSU Institutional Biosafety Committee (protocol # 2020R00000026).

### BCoV Propagation in tissue culture

BCoV-Mebus was propagated according to published protocols.^13^ Specifically, MDBK cells were seeded in T-150 tissue culture flasks, and once confluent, AMEM was removed, and cells were incubated for 3 hours in minimal essential medium (MEM, Life Technology, Catalog # 11095114) supplemented with 1x MEM non-essential amino acids (Gibco) and 1x Antibiotic/Actinomycotic cocktail (Life Technology). Cells were infected with BCoV-Mebus (median tissue culture infectious dose [TCID_50_] of 0.1 to 10) and incubated at 37°C and 5% CO_2_ for one hour. One-hour post-infection, MEM containing 6.5 µg/ml pancreatin was added and the infected cells were incubated at 37°C and 5% CO_2_ for 18 hours. Infected cells were lysed with two freeze-thaw cycles of -20°C followed by centrifugation at 500 x g for 20 minutes at 4°C. BCoV-Mebus aliquots were stored at -80°C until use.

### Passenger automobile material testing

High touchpoint surfaces inside vehicles include door handles, steering wheel, gear shift knob, turn and wiper levers, buttons or touchscreens, seat upholstery and belts, armrests, grab handles and seat adjustments, seat backs, console bins, and cup holders (Supplemental Figure 1). These components comprise a wide variety of plastics, including thermoplastic olefin (TPO), nylon 6 (PA6), poly(ethylene) terephthalate (PET), and poly(vinyl) chloride (PVC). In addition, each of these particular plastic materials is formulated with a number of fillers and additives that improve properties, processing, durability and performance characteristics. One cm^2^ pieces of each material (test coupons) were placed into a single well of a 12-well dish, in triplicate. Throughout six experiments, an average of 2.7 × 10^6^ TCID_50_ units of BCoV-Mebus was spotted as a single drop onto each test coupon and dried in a laminar flow hood (<12h). Test coupons containing dried viruses were placed into a humidity-controlled incubator set to 55% relative humidity and 56°C. Surface and air temperatures were monitored using a dual-input thermocouple (Fluke). Exposure time commenced when the surface temperature reached 56°C.

### In situ heat inactivation

One square centimeter coupons of hard plastic (seat) material were placed into a single well of a 12-well dish, in triplicate, and an average of 2 × 10^6^ TCID_50_ units of BCoV-Mebus was spotted as a single drop onto each test coupon and dried in a laminar flow hood (<4h). After drying, 12-well dishes were placed into a filter-top disposable animal cage (Innovive), and the cages were placed inside a transit bus at four locations (indicated in Supplemental Figure 2). To heat the bus, a diesel-powered, indirect fired 500,000 BTU portable heater (Frost Fighter) was attached to flexible ducts that were fed inside the bus through the back door. One duct was aimed towards the front of the bus. The second duct was bifurcated inside the bus, with one duct facing upwards and one duct facing towards the back of the bus (Supplemental Figure 2). The portion of the back door opening not occupied by the ductwork was covered with insulated blankets. The front doors were closed during the experiment, as were the windows (with factory seals).

Owing to the documented importance of humidity in coronavirus inactivation,^15^ an industrial humidifier (Ideal Air, capacity 4 liters per hour) attached to a gravity-fed 20L carboy was placed inside the bus. This system was used to humidify the bus for one hour before initiation of heat treatment. Hygrometers (VWR) were placed at two locations inside the bus, one at the front and one at the back of the bus along with four thermocouples. Thermocouples and hygrometers were monitored without entering the bus. Four filter-top animal cages containing the 12-well dishes containing seat materials spotted with bovine coronavirus were placed inside the bus at the following locations: driver ‘s seat, a front passenger seat, a middle passenger seat, and a back passenger seat. Exposure time commenced when the surface temperature reached 56°C. Control (unheated) plates remained outside of the bus at ambient temperature in a sealed container.

### TCID_50_ assay

Infectious virus was recovered from the materials by adding 1 mL of MEM followed by orbital shaking for 2 × 10 minutes (ten minutes on each side of the test coupon). After agitation, MEM was aspirated from the material and virus infectivity was measured using a TCID_50_ assay.^16^ Specifically, two days before virus infection, 10,000 MDBK cells were plated in AMEM into each well of a flat-bottomed 96-well dish. Three hours before infection, AMEM was removed and MEM supplemented with non-essential amino acids (Gibco) and antibiotic/antimycotic was added and the cells were incubated at 37°C and 5% CO_2_. Following heat treatment (or not in the case of controls), recovered virions were serially diluted (1:7) and 60 µL of each dilution was added in duplicate to confluent MDBK cells in MEM. Thus, each treated or control material was assayed in sextuplicate (triplicates of the material and duplicates of the aspirates). One-hour post-infection, an additional 60µL MEM containing 6.5 µg/ml pancreatin (Sigma) was added to the cells, and cells were incubated at 37°C and 5% CO_2_ for 48h. Cells were imaged with a SpectraMax Imaging Cytometer (Molecular Devices) to manually score cytopathogenicity (CPE). TCID_50_ values were calculated using the Reed-Muench method.^17^ Bovine Coronavirus half life in solution was calculated with GraphPad Prism version 9 using a one phase decay fit with a least squares regression.

### Phylogenetic tree construction

The placement of the BCoV-Mebus strain within the phylogeny of *Orthocoronavirinae* (including alpha, beta, gamma, and delta coronaviruses) was based on the most comprehensive evolutionary study of this virus subfamily to date.^18^ The phylogenetic tree was constructed from 2,006 complete and unique genomes of coronaviruses (12 deltacoronavirus, 265 gammacoronaviruses, 630 alphacoronaviruses, and 1,099 betacoronaviruses) downloaded from NCBI and GISAID. In order to avoid common errors that were identified in leading publications of the evolution of viruses,^19,20^ we used several techniques including: 1) successive outgroup expansion,^21^ 2) genome annotation and multiple sequence alignment (using MAFFT v7.453^22,23^) of homologous gene partitions (for the polyprotein 1ab, spike, membrane, envelope, and nucleoprotein genes). Heuristic searches were conducted under the parsimony (using TNT v1.1^24^) and maximum likelihood (using IQ-TREE v1.6.12^25–28^) optimality criteria. Additionally, we addressed the potential impact of putative recombination events in a subset of 505 terminals using RDP v5^29^ with the RND and GENECOV algorithms. Additional details on the data and methodological procedures used for phylogenetic analyses have been previously described.^30^

## Results

### Selection of a Surrogate Coronavirus

Worldwide, Bovine coronavirus is a causative agent of diarrhea in newborn calves, winter dysentery in adult cattle, and respiratory tract illnesses.^31^ The disease leads to significant economic losses in the beef and dairy industry.^32^ Bovine coronaviruses and bovine-like coronaviruses also have well characterized genomes and genetic features.^33,34^ Standard reference strains of BCoV include the Quebec and Mebus (GenBank accession numbers AF220295.1 and U00735.2, respectively) strains.^35^ The most recent and comprehensive phylogenetic analysis of coronaviruses^18^ counts with 2,006 unique genomes of all *Orthocoronavirinae* genera (*Deltacoronavirus, Gammacoronavirus, Alphacoronavirus*, and *Betacoronavirus*). That analysis unequivocally places BCoV strain Mebus within a clade that includes human coronaviruses (HCoVs) such as HCoV-HKU1, HCoV-0C43, and HCoV-4408 (Figure 1). The phylogenetic position of BCoV strain Mebus favors its strategic application as a safe proxy to Human CoVs (HCoV). The BCoV strain Mebus is phylogenetically related to HCoVs associated with mild human diseases (see the blue group in Figure 1). This relation to HCoVs indicates Mebus is a potential proxy to betacoronaviruses of clinical importance in controlled experiments. Simultaneously, BCoV strain Mebus is placed outside from groups in which host transformations from other animals to humans led to severe diseases, including COVID-19 (see the red group in Figure 1). Therefore, BCoV strain Mebus is sufficiently distant from the SARS-CoV, MERS-CoV, and SARS-CoV-2 strains that it can be safely used in experiments where human contact could occur.

**Figure 1.**
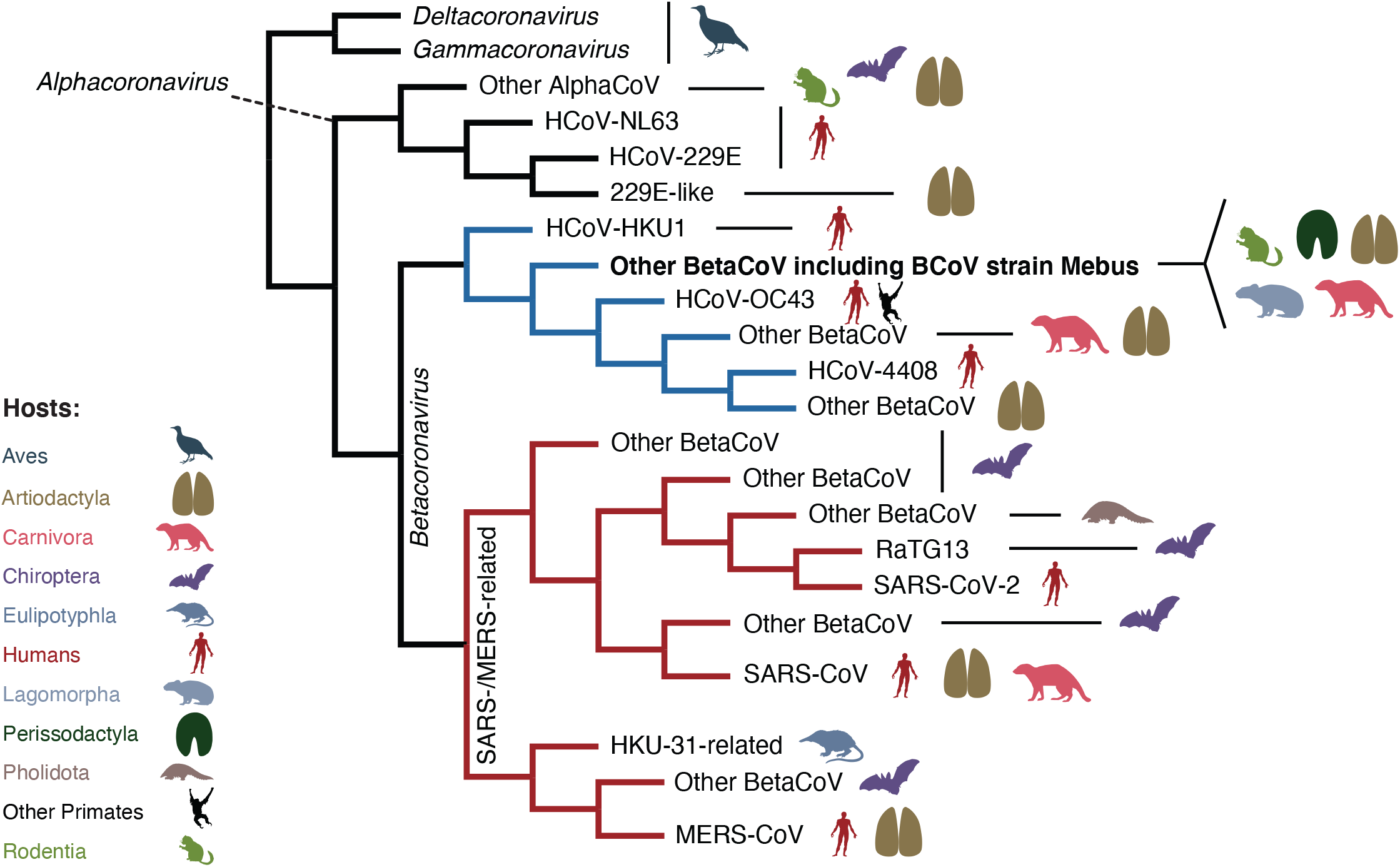
Simplified coronavirus consensus tree. The best heuristic results from parsimony analysis were congruent with the maximum likelihood tree. The terminal in bold represents the Bovine coronavirus (BCoV) strain Mebus and other betacoronaviruses, nested within a clade that includes human coronaviruses (HCoVs) known to cause mild disease in humans (indicated in blue). The group in red represents lineages of coronaviruses known to cause severe diseases in humans (SARS, MERS, and COVID-19). Redrawn from Fig. S1.1 in Machado et al., (2021).^18^

### Heat-based inactivation of coronaviruses

Having identified BCoV-Mebus as an accessible surrogate coronavirus, we next tested the ability of heat to inactivate the virus. Others have shown that passive solar heating of vehicles can achieve air temperatures of 56°C.^9^ Ford Motor Company has demonstrated a unique software solution that controls the powertrain and climate control systems that can be used to increase and hold interior cabin surface temperatures to 56°C.^36^ To quantify the effect of 56°C on BCoV infectivity, we incubated a solution of 3.64 × 10^5^ ± 2 × 10^5^ (n=4) TCID_50_ units of BCoV in MEM at 56°C for 0, 15, 20, 30, or 60 minutes. In comparison to the unheated virus, heating the virus suspension at 56°C for 15 minutes reduced BCoV infectivity by over 1000-fold. We were unable to recover any infectious BCoV following heating at 56°C for 30 or 60 minutes (Figure 2). Fitting the data to a single-phase decay returned an average half-life at 56°C of 1.3 minutes (95% Confidence Interval: 1.0, 1.4).

**Figure 2:**
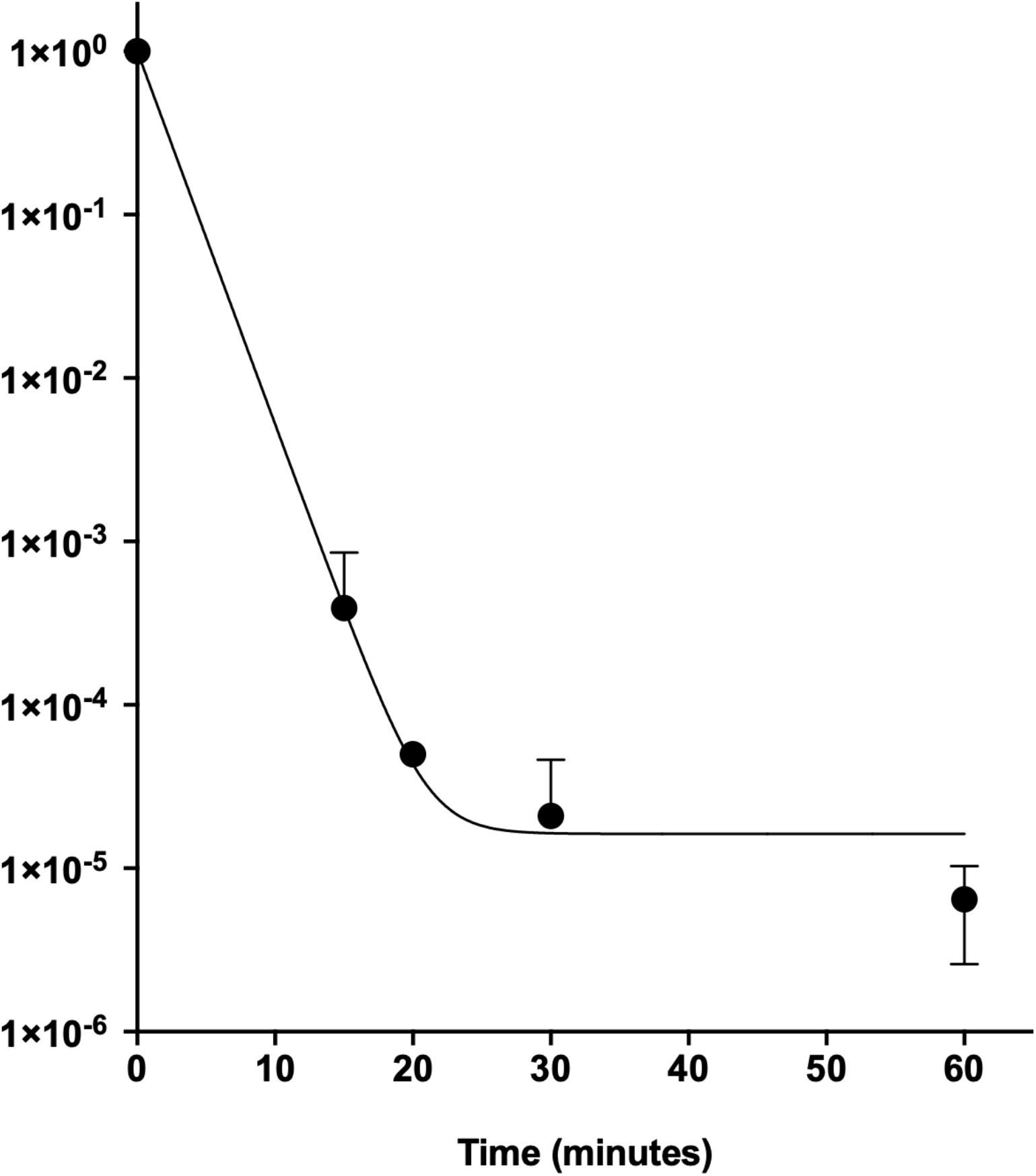
Bovine coronavirus (BCoV) decay at 56°C. BCoV (strain Mebus) suspended in MEM was placed into a PCR tube and heated at 56°C for the indicated times (n=4). Unheated virus, heated virus, and no virus control reactions were incubated with MDBK cells in sextuplicate for 48 hours, cytopathogenecity (CPE) was scored, and TCID_50_ values were calculated. TCID50 values for each experiment were normalized to the unheated (time = 0) time point. The limit of detection (LoD) for the CPE assay is equivalent to TCID_50_=1. When CPE was undetectable, TCID_50_ was set to 1 (LoD). Point estimate, standard deviation, and best fit line for a single-phase exponential decay are plotted. The half-life of BCoV at 56°C is 1.3 minutes (95% confidence interval = 1.0, 1.4).

Next, we obtained swatches of a representative group of porous and non-porous materials commonly found inside passenger vehicles (on touch available surfaces) and placed a single drop of BCoV onto them (Table 1); virus-laden materials were placed inside a biosafety cabinet until dry. Materials containing dried BCoV were heated in an incubator at constant humidity until the surface temperature achieved 56°C (average time to achieve 56°C surface temperature = 29 ± 7 minutes, n=6). Next, the virus-laden materials were held at 56°C for 15 minutes. In all experiments, infectious virus was never recovered from materials heated to 56°C for 15 minutes, resulting in 99.99% to 99.9999% reduction in virus infectivity when compared to the virus stock solution. To calculate the relative reduction in viral infectivity, we compared the TCID_50_ of BCoV recovered from untreated materials to the TCID_50_ of BCoV recovered from 56°C-treated materials. Compared to the untreated group, a surface temperature of 56°C held for 15 minutes was sufficient to inactivate BCoV spotted on all materials, equivalent to a >99% to 99.99% reduction in the amount of infectious virus recovered (Table 1). Recovery of infectious virus from untreated materials was highly variable, and because reduction in virus infectivity depends on recovery of infectious virus from the untreated materials, the magnitude of reduction of infectivity was influenced by virus interactions with and adherence to the materials.

**Table 1:** Inactivation of Bovine Coronavirus on materials found inside automobile cabins. A single spot of Bovine coronavirus (Mebus) was dried onto several typical automotive surface materials, heated to a surface temperature of 56°C (50% relative humidity), and held at 56°C for fifteen minutes. Infectious bovine coronavirus (mebus) was recovered from both heated and unheated materials and cytopathic effects (CPE) were quantified by visual scoring 48h post-infection. Nd: below the level of detection; PVC: polyvinyl chloride; PET: polyethylene terephthalate; PA6: Polyamide; TPO: thermoplastic olefin.

| MATERIAL | TCID<sub>50</sub><br>(unheated) | TCID<sub>50</sub><br>(heated) | Reduction |
| --- | --- | --- | --- |
| Virus suspension (liquid) | 1.98E+06 | N/A | N/A |
| Tissue culture dish plastic | 3.02E+04 | nd | >99.99% |
| PVC black vinyl, seat and trim covering | 5.50E+02 | nd | >99% |
| PET for automotive interior fabric | 1.55E+02 | nd | >99% |
| PET stamped fabric | 7.84E+02 | nd | >99% |
| (PET or nylon 6) carpet floor mats | 1.12E+02 | nd | >99% |
| 30% glass fiberfilled nylon6, mold-in-color for door handle applications | 5.49E+03 | nd | >99.9% |
| 20% talc-filled thermoplastic olefin (TPO) (medium impact and high scratch) for class-A, visible interior surfaces | 5.48E+04 | nd | >99.99% |
| PVC Floor vinyl | 4.06E+04 | nd | >99.99% |

### Heat-based coronavirus inactivation in a transit bus

To test the ability of heat to inactivate BCoV *in situ*, we spotted BCoV onto plastic used to form the seats of a public transportation bus. The plastic coupons containing dried BCoV were placed inside of 12 well tissue culture dishes placed inside disposable, filter-topped plastic cages, which were then placed at four locations inside the bus. Next, the bus doors were closed and the humidified vehicle was heated with a portable, diesel powered heater until surfaces inside the bus reached a temperature of 56°C (approximately 75 minutes); once a surface temperature of 56°C was achieved, the virus-laden samples remained on the bus for an additional 30-minute incubation (heat and humidity profiles plotted in Supplemental Figure 2B). To calculate the relative reduction in viral infectivity, we compared the TCID_50_ of BCoV recovered from unheated (ambient temperature) materials to the TCID_50_ of BCoV recovered from the heated materials. Recoverable virus infectivity from materials placed on the driver ‘s seat and the front of the bus was reduced by 99.5%, while the middle and back of the bus was reduced by 85% (Table 2).

**Table 2:** Inactivation of Bovine Coronavirus inside the cabin of a transit bus. A single spot of Bovine coronavirus (Mebus) was dried onto plastic used to form seats, virus-laden coupons of the seat plastic were placed throughout the bus (Supplemental figure 2), the interior of the bus was humidified and heated to a surface temperature of 56°C, and held at 56°C for thirty minutes. Infectious bovine coronavirus (mebus) was recovered from both heated and unheated material and cytopathic effects (CPE) were quantified by visual scoring 48h post-infection. Nd: below the level of detection.

| Location | TCID<sub>50</sub> | Reduction in infectivity following incubation at 56°C for 30 minutes |
| --- | --- | --- |
| Virus suspension (liquid) | 2.23E+06 | N/A |
| No heat control | 5.06E+02 | Referent |
| Driver | nd | 99.5% |
| Front | nd | 99.5% |
| Middle | 7.73E+01 | 85% |
| Back | 7.56E+01 | 85% |

## Discussion

Like several other viruses, coronaviruses can be inactivated by exposure to heat in a manner dependent on the matrix, humidity, and temperature.^37^ We observed that BCoV in suspension in MEM had a half-life of 1.3 minutes when heated to 56°C (Figure 1), which is similar, albeit it shorter, than the reported half-life of SARS-CoV-2 in a matrix of artificial saliva (half-life of 10.8 minutes at 56°C^15^). Using swatches of a variety of materials found inside the interior cabin of an automobile, we observed that heating the surface of these materials to 56°C followed by a 15-minute hold was sufficient to inactivate 99% - 99.99% of infectious viruses. In a real-world, in situ test of a transit bus warmed with an external heater, a surface temperature of 56°C followed by a 30-minute hold reduced infectivity of BCoV by 85% - 99.5%.

Several sanitation methods can inactivate viruses, including chemical disinfectants,^38^ ultraviolet light,^39^ and heat. According to the US Environmental Protection Agency (EPA) guidance on cleaning and disinfecting public spaces, a primary means to disinfect public and private enclosed spaces comprises disinfecting surfaces with EPA-approved products (List N) or a bleach solution or a 70% alcohol solution. The wipe down method is effective if the solution is applied to the contaminated surfaces according to label instructions, typically for an extended period before being wiped clean. While effective, there are several limitations to wiping down contaminated surfaces, including the following: 1) wiping down is laborious, especially when dealing with a fleet of vehicles, 2) it can be difficult to determine if the disinfectant contacts all surfaces in a complex space, and 3) solutions that contain bleach, hydrogen peroxide and ammonia can damage, discolor, and weaken materials.^40^ Similar limitations are seen with ultraviolet light, which can be effective with direct contact but suffers from shadowing and penetration issues when treating complex surface structures, crevices, and course fabrics.^39,41,42^

Alternatively, heat can penetrate materials, is not sensitive to shadowing, is easy and inexpensive to generate, and is well tolerated (to a point) by materials at levels that inactivate viruses.^43^ For instance, long term heat durability testing on vehicles is already performed at temperatures up to 80°C, as air temperatures inside of a closed vehicle can exceed 90°C on a hot day. Therefore, the polymer materials used in automotive are already stable at 56°C for long exposures. If for some instance they are not, well-known heat stabilizers called hindered amine light stabiliizers (HALS) can be added to extend heat performance by reducing free radical degradation mechanisms. Heat works to inactivate viruses by denaturing secondary structures of viral proteins^44^ and other biomolecules, rendering the virus particle non-infectious.^45^ Although there are many advantages to using heat as an inactivation agent, there are also some limitations to consider. Heating to the desired surface temperature (56°C) could take longer than wiping down surfaces, and there are likely to be micro climates inside of the heated spaces, which may not achieve the heat and humidity^49^ parameters required for efficient virus inactivation. We attempted to minimize microclimates inside the bus by circulating the air with fans, but the observed variability in virus inactivation suggests that the additional optimization of heat and humidity conditions are required to achieve optimal conditions for virus inactivation. Nevertheless, in the context of automobile sanitization, heat is readily available through passive heating, through software-solutions that alter the normal function of the powertrain and the climate control system, or via portable hot air blowers. Other potential sources include high temperature heat systems currently used for the remediation of Bed Bugs (*Cimex lectularius*) throughout the pest control industry. Most of these systems can easily reach and maintain the desiredinactivation temperatures of 56°C. Importantly, heat permeates structures and also sanitizes the air, so long as the air and surfaces achieve the desired temperature and are held for the determined amount of time. Moreover, unlike the wipe-down method, heat will also inactivate aerosolized viral particles,^46^ which contribute to the transmission of SARS-CoV-2.^47,48^ Thus, heat appears to be a simple and superior solution, overcoming several limitations of wipe-down, UV light and other sanitation methods.

In conclusion, our in vitro and in situ results indicate that 56°C heat is an efficient, inexpensive, and effective method to disinfect virus laden touchpoints on common vehicle materials, and given that heat is a practical, safe, and widely applicable, it should be considered as a mode of virus inactivation for enclosed vehicle spaces.

## Acknowledgements

This work was supported by the OSU Infectious Diseases Institute, Ford Motor Company Research and Innovation Center, and the Cooper Family Foundation. DJM and DJ acknowledge support from UNC Charlotte and the Belk family. We thank Covectex, LLC for the provision of heaters for optimization of the in situ experiments, the OSU Center for Automotive Research (CAR) for logistical support of the transit bus experiments, and Central Ohio Transit Authority (COTA) for providing an out of service transit bus and seating material samples.

