## Supplementary Figures for "Heat efficiently inactivates coronaviruses inside vehicles"

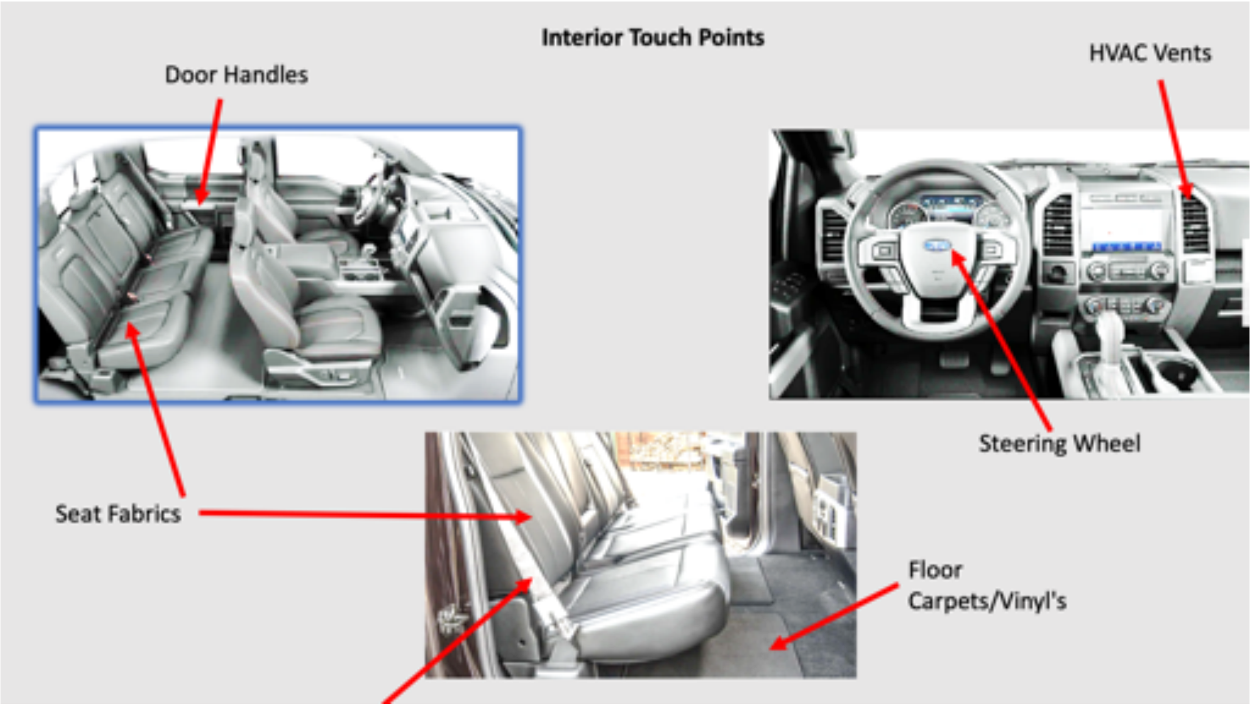


**Supplemental Figure 1. Interior touch points.** High touch surfaces were identified and representative materials from these surfaces were selected for laboratory-based studies to quantify the ability of 56°C heat to inactivate bovine coronaviruses.

A.

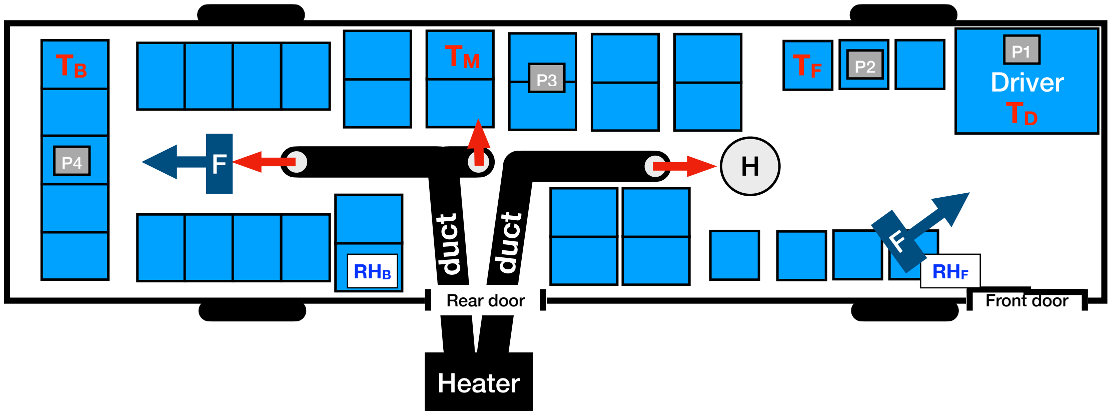


B.

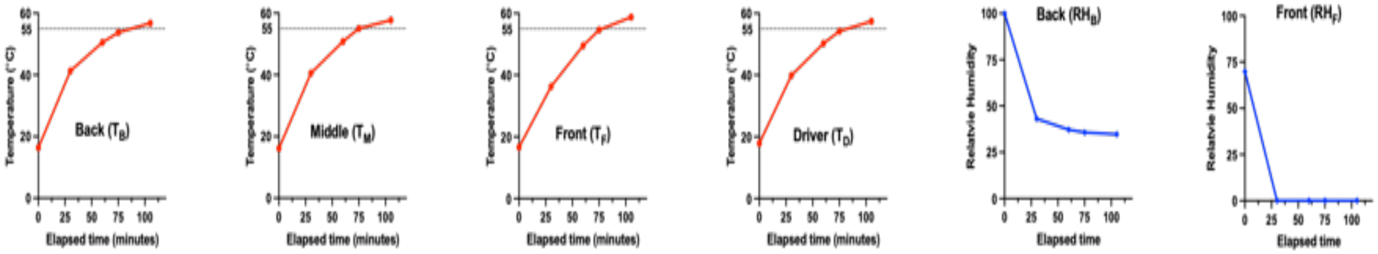


**Supplemental Figure 2: Transit bus layout, temperature, and humidity profiles.** A). An out of service public transit vehicle was heated until interior surfaces reached 56°C. Diesel powered heater (H) was placed outside the bus and the head was transferred to the bus with two insulated ducts (in black). Four plastic cages (P1 - P4) containing dried coronavirus spotted onto seat material were placed on seats (blue) throughout the bus. Air was humidified with a commercial humidifier (H) and circulated with several fans (F). B) Hygrometers ($RH_F$ & $RH_R$) and temperature probes ($T_D$, $T_F$, $T_M$,$T_B$,) were placed throughout the bus and temperature and humidity were measured over the course of the experiment.

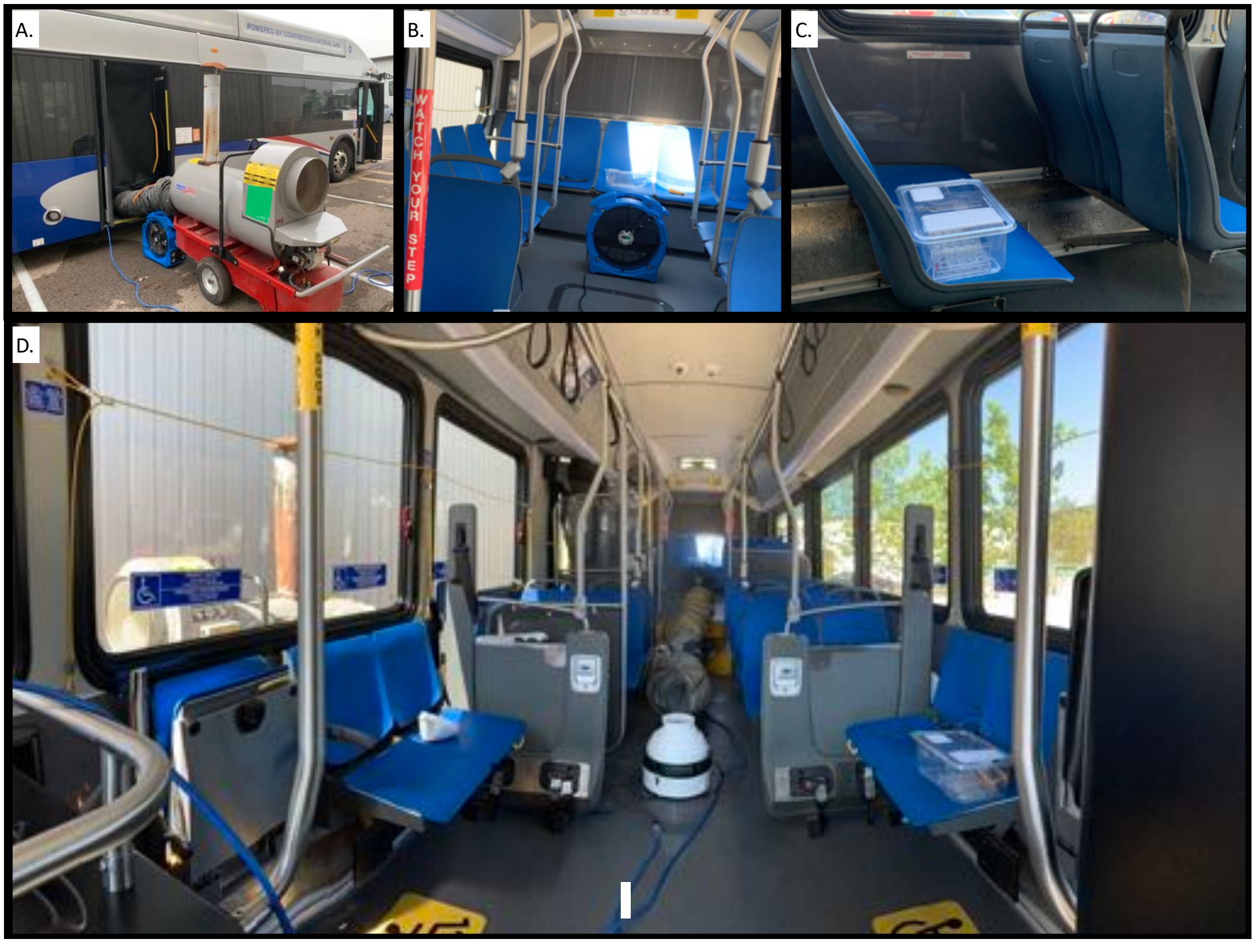

**Supplemental Figure 3: Photographs of the in situ testing setup.** A) Heater outside of bus with ducts entering the covered rear door, B) Fan and test plate in the back of the bus, C) test plate situated in the middle of the bus, D) view of the inside of the bus from front of the bus.
